# A benchmark of gene expression tissue-specificity metrics

**DOI:** 10.1101/027755

**Authors:** Nadezda Kryuchkova-Mostacci, Marc Robinson-Rechavi

## Abstract

One of the major properties of genes is their expression pattern. Notably, genes are often classified as tissue-specific or housekeeping. This property is of interest to molecular evolution as an explanatory factor of, e.g., evolutionary rate, as well as a functional feature which may in itself evolve. While many different methods of measuring tissue specificity have been proposed and used for such studies, there has been no comparison or benchmarking of these methods to our knowledge, and little justification of their use. In this study we compare nine measures of tissue-specificity. Most methods were established for ESTs and microarrays, and several were later adapted to RNA-seq. We analyze their capacity to distinguish gene categories, their robustness to the choice and number of tissues used, and their capture of evolutionary conservation signal.

## Introduction

Gene expression analysis is widely used in genomics, measured with microarrays or RNA-seq. In the case of a multicellular organism with different tissues, it is often useful to have a measure of how tissue-specific a gene is.

Even if tissue specificity is often used in studies, there is usually no clear answer why one or another method was used. Yet there are several methods to measure gene specificity, which differ in their assumptions and their scale. The simplest one is to count in how many tissues each gene is expressed (used in e.g. (Subramanian & Kumar 2004; Duret & Mouchiroud 2000; Park & Choi 2010; Lercher et al. 2002; Vinogradov 2003; Ponger et al. 2001)). The problem of this method is to define the threshold to call a gene expressed. Originally, with ESTs a count of 1 EST was considered sufficient (Duret & Mouchiroud 2000). There are different methods to define thresholds for microarrays (Liu et al. 2002), while for RNA-seq an RPKM value of 1 is generally used (Wagner et al. 2013; Hebenstreit et al. 2011). Some studies use a very stringent threshold, e.g. signal to noise ratio greater then 10 (Dezso et al. 2008), and count a gene as specific only if expressed in a single tissue. This method causes only highly expressed genes to be taken into account; and if a data set contains closely related tissues (e.g., brain parts), less genes are called tissue specific. Other papers use a very low threshold, e.g. 0.3 RPKM (Ramsköld et al. 2009; Ma et al. 2014; Cui et al. 2012), that leads to defining most genes as housekeeping.

A widely used method which does not depend on such a cut-off is Tau (Yanai et al. 2005). Tau varies from 0 to 1, where 0 means broadly expressed, and 1 is very specific (used in e.g. (Smeds et al. 2015; Piasecka et al. 2012; Assis & Bachtrog 2013; Assis & Kondrashov 2014; Bush et al. 2015; Kryuchkova-Mostacci & Robinson-Rechavi 2015; Liao & Zhang 2006; Liao et al. 2006; Weber & Hurst 2011; Zhao et al. 2015)).

Other methods have been proposed, such as the expression enrichment (EE) (Yu et al. 2006), used to calculate for which tissue each gene is specific, for example in the database TiGER (Liu et al. 2008). We also considered: the tissue specificity index TSI (Julien et al. 2012) (used in e.g. (Cortez et al. 2014; Assis & Bachtrog 2015; Winter et al. 2004)); Hg by Schug et al. (Schug et al. 2005); the z-score (used in (Vandenbon & Nakai 2010)), widely used for other features than tissue specificity; SPM, used in the database TiSGeD (Xiao et al. 2010); PEM, Preferential Expression Measure, suggested for ESTs by Huminiecki et al. (Huminiecki et al. 2003) and used in e.g. (Milnthorpe & Soloviev 2012; Russ & Futschik 2010; Lin et al. 2008; Divina et al. 2005). Finally, the Gini coefficient, widely used in economics to measure inequality (Ceriani & Verme 2012), was compared to methods originating in biology.

These methods can be divided in two groups. One group summarizes in a single number whether a gene is tissue specific or ubiquitously expressed (Tau, Gini, TSI, Counts, Hg), and the second group shows for each tissue separately how specific the gene is to that tissue (Z score, SPM, EE, PEM). For comparison purposes with the first group, we use the maximum specificity from the second group.

## Material and methods

For all equations apply:

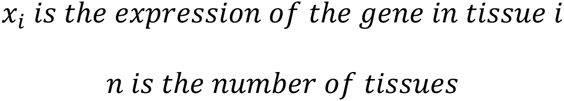

The method of counting in how many tissues a gene is expressed was simply calculated as follows:

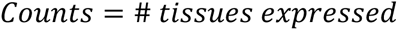

A cut-off needs to be set; the cut-offs that we used are explained at the end of the Methods section.

Tau was calculated as follows (Yanai et al. 2005):

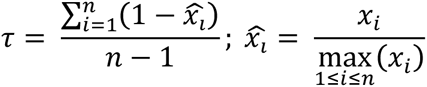

The EE score was calculated as follows (Yu et al. 2006):

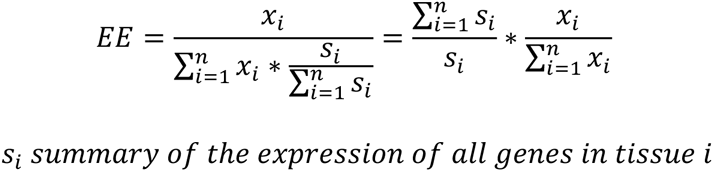

TSI was calculated as follows (Julien et al. 2012):

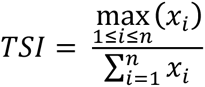

The Gini coefficient was calculated as follows:

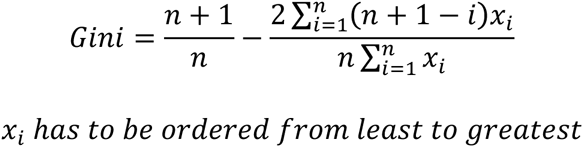

Hg (Schug et al. 2005) was calculated as follows:

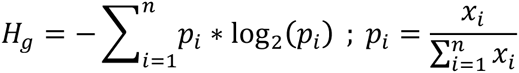

The z-score was calculated as follows:

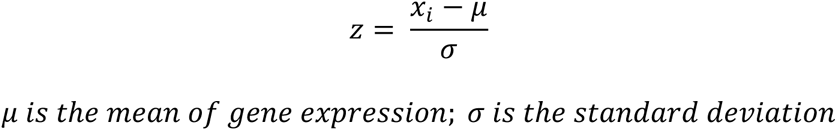

The z-score can be implemented in two ways: either only over expressed genes are defined as tissue specific, or the absolute distance from the mean is used, so that under-expressed genes are also defined as tissue specific. Only the former method was used to be able to compare z-score to other methods.

SPM from the database TiSGeD (Xiao et al. 2010) was calculated as follows:

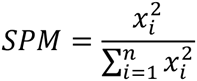

PEM estimates how different the expression of the gene is relative to an expected expression, under the assumption of uniform expression in all tissues. PEM is calculated as follows (Huminiecki et al. 2003):

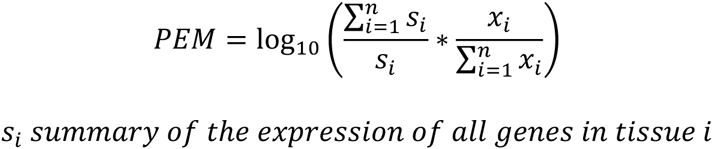

Derivation of all the methods from the original equations is presented in Supplementary Materials.

The output of all methods was modified to the same scale from 0 (ubiquitous) to 1 (tissue specific) to be able to compare them (Table 1). Four of the methods calculate specificity value for each tissue separately; for these methods the largest (most specific) value among all tissues was assigned to the gene (see Table 1).

All the methods were compared using R version 3.2.1 (R Core Team 2015), with the gplots (Warnes et al. 2015), reldist (Handcock & Morris 1999; Handcock 2015), VennDiagram (Chen 2015) and preprocessCore libraries (Bolstad); the R script is available in Supplementary Materials.

**Table 1:**
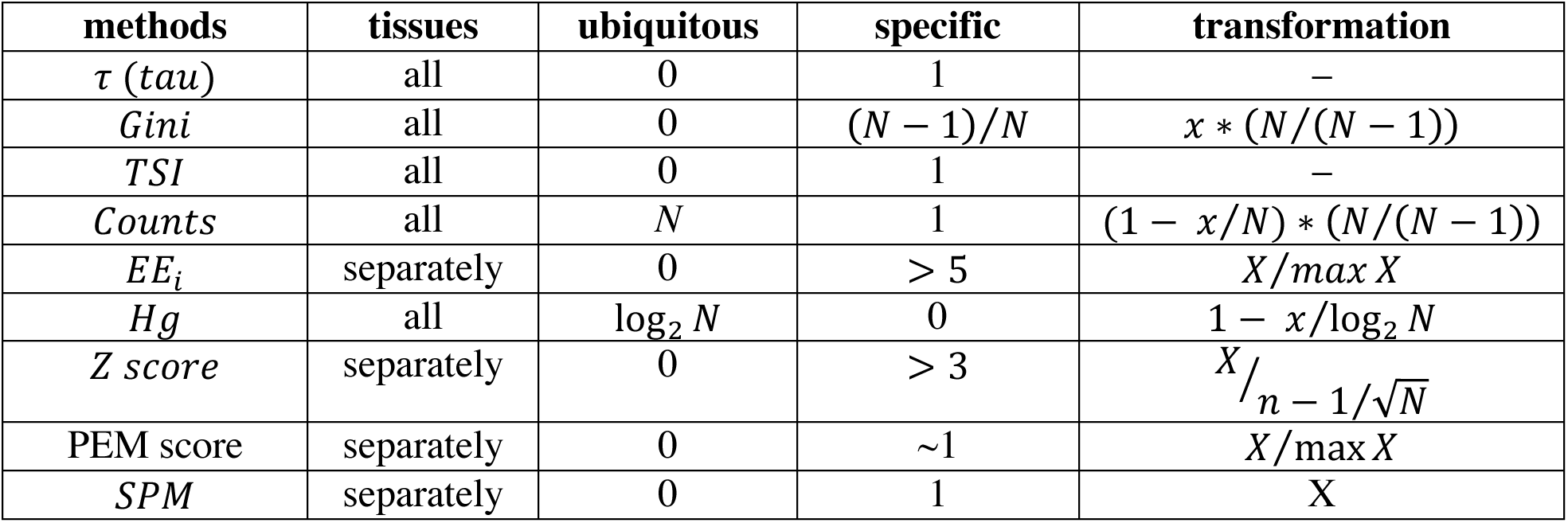
Tissue specificity parameters. *N* is the number of tissues in the data set. *X* = max_1<_*_i_*_<_*_N_ x_i_* is the maximal specificity value for a certain gene among all tissues.

We used the following RNA-seq data: 27 human tissues (E-MTAB-1733) from Fagerberg et al. (Fagerberg et al. 2013) downloaded from their Supplementary Materials, 22 mouse tissues (GSE36025) from the ENCODE project (The ENCODE Project Consortium 2011; Lin et al. 2014) as used in Kryuchkova-Mostacci & Robinson-Rechavi (Kryuchkova-Mostacci & Robinson-Rechavi 2015), and 8 human tissues and 6 mouse tissues from Brawand et al. (Brawand et al. 2011), as processed in the Bgee database (Bastian et al. 2008). All the genes with expression less than 1 RPKM were set as not expressed. The RNA-seq data were first log-transformed. After the normalization a mean value from all replicates for each tissue separately was calculated. All genes that were not expressed in at least one tissue were removed from the analysis.

We used the following microarray data, as annotated in the Bgee database: 32 human tissues (GSE2361) (Ge et al. 2005), and 19 mouse tissues (GSE9954) (Thorrez et al. 2008). Of note, on the microarrays we have only 9788 (resp. 16043) genes with data in human (resp. mouse), relative to 18754 (resp. 27364) for RNA-seq. For the microarray data we used the logarithm of normalized signal intensity. The values set as absent in Bgee were set to 0, following the method of Schuster et al. (Schuster et al. 2007). After the normalization a mean value from all replicates for each tissue separately was calculated. All genes that were not expressed in at least one tissue were removed from the analysis.

A summary of the workflow is presented in supplementary Figure S1.

For the comparison of tissue-specific or ubiquitous gene functions, we used the following Gene Ontology terms: spermatogenesis (GO:0007283; expected to be specific to testis; 469 human genes), neurological system process (GO:0050877; expected to be specific to brain and other neural tissues; 1338 human genes), xenobiotic metabolic process (GO:0006805; expected to be specific to liver and kidney; 163 human genes), protein folding (GO:0006457; expected to be ubiquitous; 231 human genes), membrane organization (GO:0061024; expected to be ubiquitous; 607 human genes), and RNA splicing (GO:0008380; expected to be ubiquitous; 383 human genes).

Gene Ontology enrichment analysis was performed with GOrilla (Eden et al. 2009) and Revigo (Supek et al. 2011).

## Results

All methods show a bimodal distribution of gene scores: most genes are either broadly expressed or specific, with only few in between. This is true both with RNA-seq data (Figure 1 and S2) and with microarray data (Figure S3 and S4). Most methods are strongly skewed towards classifying many genes ubiquitous, and very few tissue-specific or intermediate. Z-score has a shifted peak of tissue-specificity relative to other metrics. Tau has a less skewed distribution, with the most tissue-specific and intermediate genes, indicating that it might be capturing more of the variance among gene expression patterns.

All methods correlate relatively well with each other (Figure S5 and S6), but the relation is often not linear because other methods than Tau and Gini have very little variance outside of the extreme most tissue-specific genes. For example, genes which have Tau between 0.85 and 0.95 have Tsi between 0.2 and 0.43 (Figure S5).

**Fig. 1:**
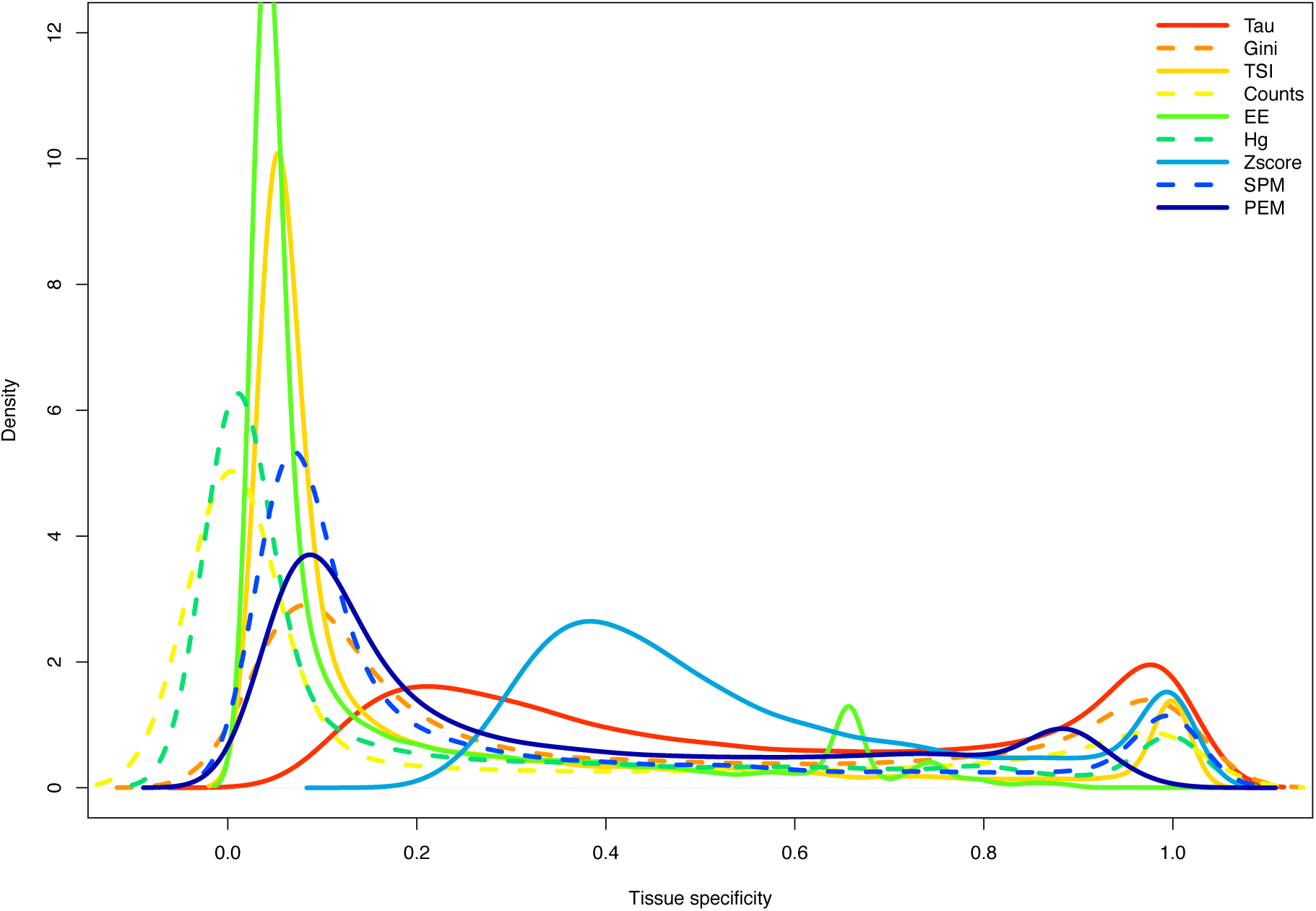
Distribution of tissue specificity parameters with data for human RNA-seq 27 tissues. Graph created with density function from R, which computes kernel density estimates.

As a first measure of robustness of tissue-specificity metrics, we compared each metric calculated on the full human RNA-seq data set of 27 tissues, and on subsets of 5 tissues (Figure 2). Not all permutations were performed, for computational reasons, but a random sample of 1000 permutations. Ideally, the signal for tissue-specificity should already be detectable with the 5 tissues. Tau, Gini, Counts, PEM and the Hg coefficient all show correlations which are not too low (mean r > 0.4), indicating that these methods are reasonably robust to the number of tissues. TSI, SPM and EE score show weaker results (mean 0.2 > r > 0). The correlation for z-score is even negative, indicating that it should be not used with a small number of tissues, and casting doubt on its utility to robustly estimate tissue-specificity. We performed the same analysis in mouse, comparing scores between all 22 available tissues and subsets of 5 tissues; the results are consistent but correlations are weaker for all parameters (Figure S7). Similarly, we compared the scores using all available tissues (27 in human, 22 in mouse) to the scores using only the 16 tissues shared between these human and mouse datasets; correlations of all parameters are high for human and mouse, and z-score shows again the lowest correlation in all cases (Figure S8 and S9).

The choice of tissues to calculate tissue specificity affects the results. All the outliers (stronger correlation) in Figure 2 and S7 contain testis. This can be explained by the fact that testis has the largest number of tissue specific genes (Figure S10 and S11). Thus using a subset which excludes testis produces an estimate of tissue-specificity which is biased relative to the full dataset, and this bias is only relieved in the few subsets which include testis.

We also analysed robustness of Tau by comparing correlation calculated on all 27 tissues and on all the subsets of 5 to 26 tissues (Figure S12 and S13). Again, all the subsets which are most similar to the full set (outliers with r > 0.7) in subsets of 5 and 6 tissues contain testis in the set. Conversely, all the subsets which are most different in the full set (outliers with r < 0.8) in subsets of 21 to 26 tissues do not have testis in the subset. There are other outlier subsets which are closer to the main distribution for 25 or 26 tissues: these do include testis, but not brain, which is the second tissue with the most specific genes.

**Fig. 2:**
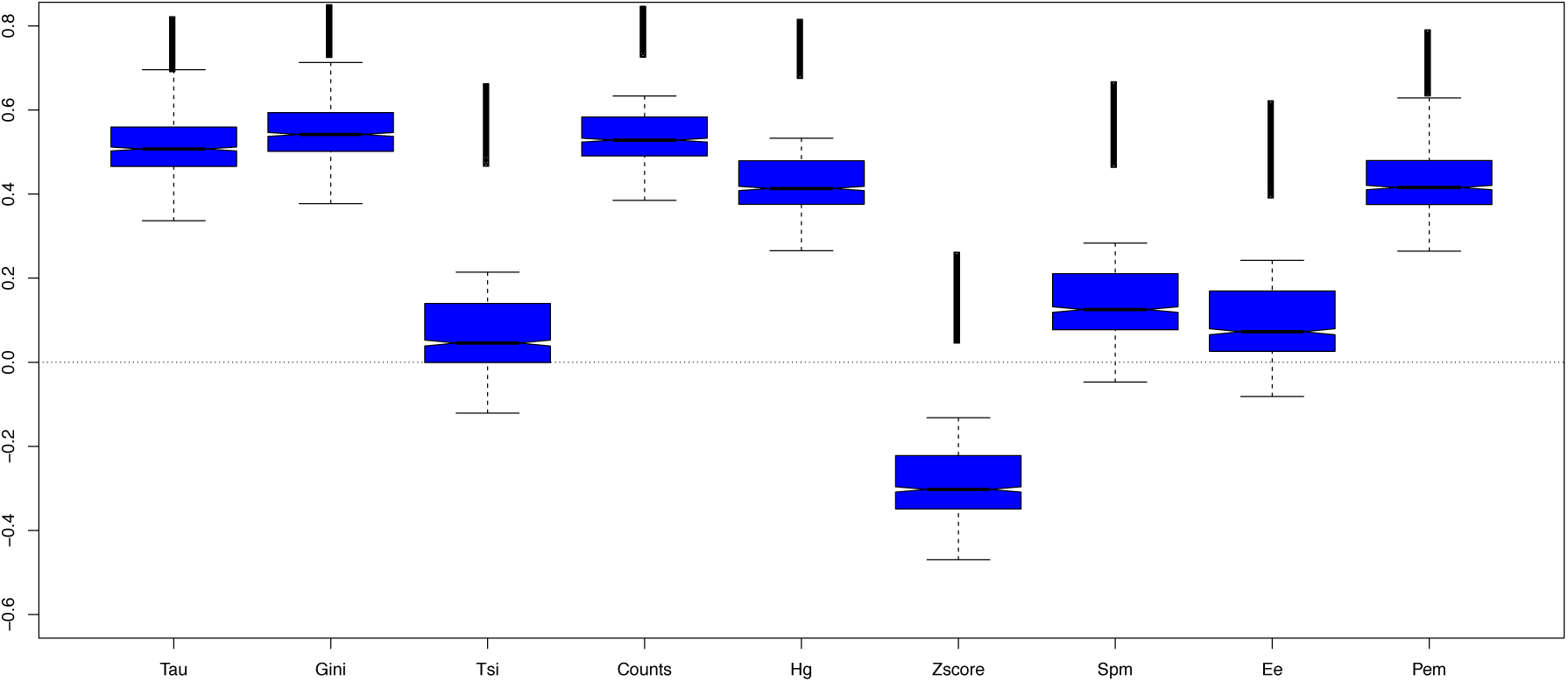
Comparison between tissue specificity parameters calculated on the same human RNA-seq data set using all 27 tissues vs. 1000 random subsets of 5 tissues.

In addition to being robust to tissue sampling, we expect a good measure of tissue-specificity to capture biological signal. A simple expectation of such biological signal is that it should be mostly conserved between orthologs from closely related species such as human and mouse (Rosikiewicz & Robinson-Rechavi 2014). Thus, we compared the methods in their conservation between human and mouse, using the 16 common tissues (Figure 3). All of the methods, except z-score, show a high correlation (r > 0.69). Specificity parameters calculated on only 6 common tissues between mouse and human (from the Brawand et al. data set) show even higher correlations (r > 0.75, Figure S14).

**Fig. 3:**
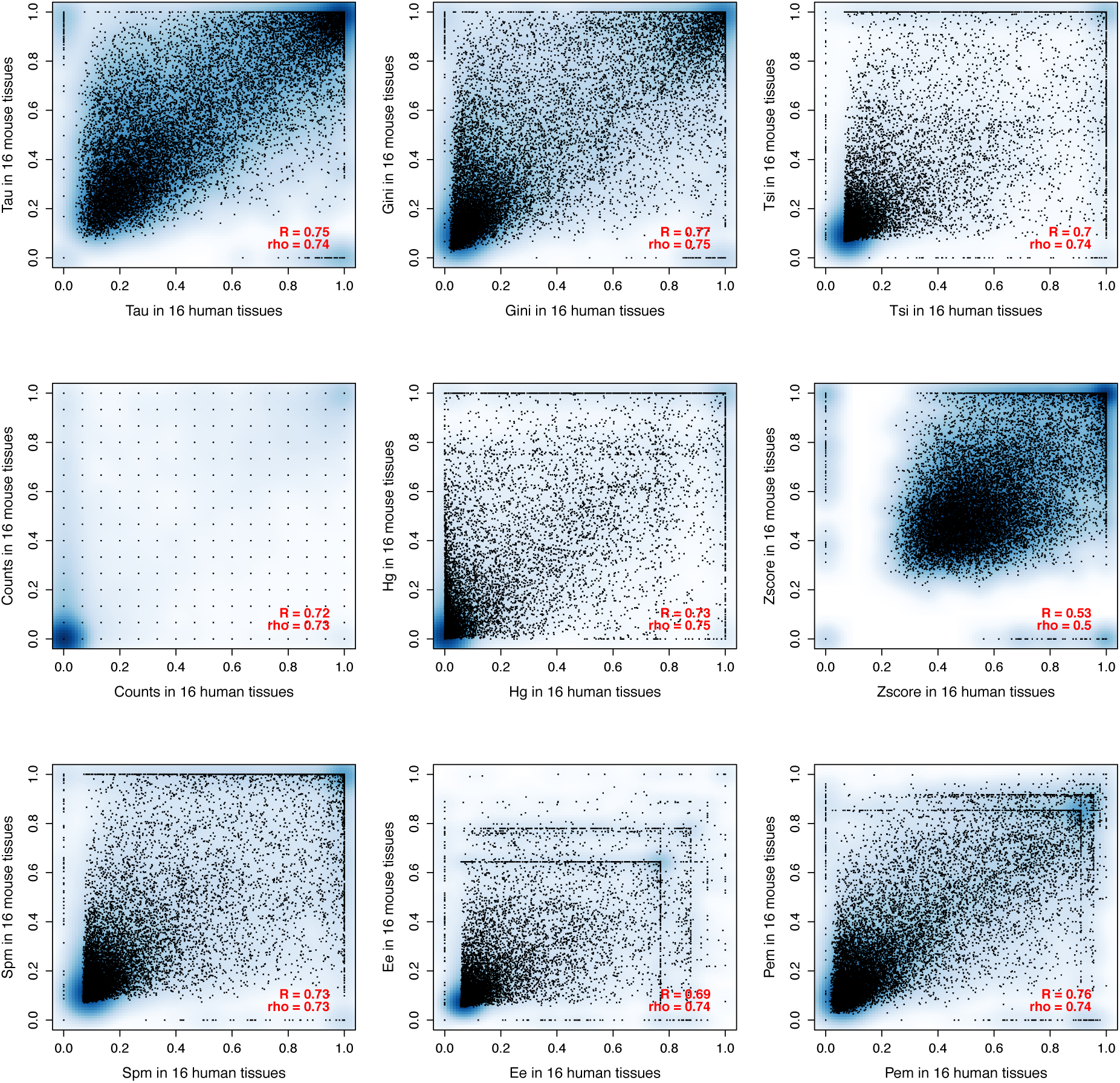
Comparison between tissue specificity parameters calculated on the 16 common tissues between the human and mouse RNA-seq data sets. All correlations have p-value < 2.2^*^10^−16^.

Another way to capture biological signal is to compare the expression-specificity of genes annotated with functions which are expected to be very tissue-specific, or which are expected to be very ubiquitous. For this we chose three tissue-specific GO terms and three GO terms that are expected to be present in all tissues. The tissue specific GO terms are spermatogenesis, specific to testis, neurological system process, specific to brain and other neural tissues, and xenobiotic metabolic process, specific to liver and kidney. The broadly expressed GO terms are protein folding, membrane organization and RNA splicing. The distribution of the genes belonging to each category is presented in Figures 4 and S15. All of the parameters are successful at recognising broadly expressed genes (peak of blue lines as expected shifted towards 0). But there are important differences in results for specific genes. Only Tau has a larger peak close to 1 than close to 0. All parameters, except Tau, show strongly bimodal distributions for the genes that are expected to be specific, often with the larger peak at ubiquitous expression. Thus Tau appears to be more successful at recovering this expected biological signal. We also checked the correlation of genes from tissue-specific functions (according to the three GO terms) between mouse and human orthologs (Figure S16). Even if correlations are high and almost the same for all the parameters, the difference is that genes that are expected to be specific are specified by most parameters as ubiquitous. Only Tau reports most of these genes as evolutionarily conserved tissue-specific.

**Fig. 4:**
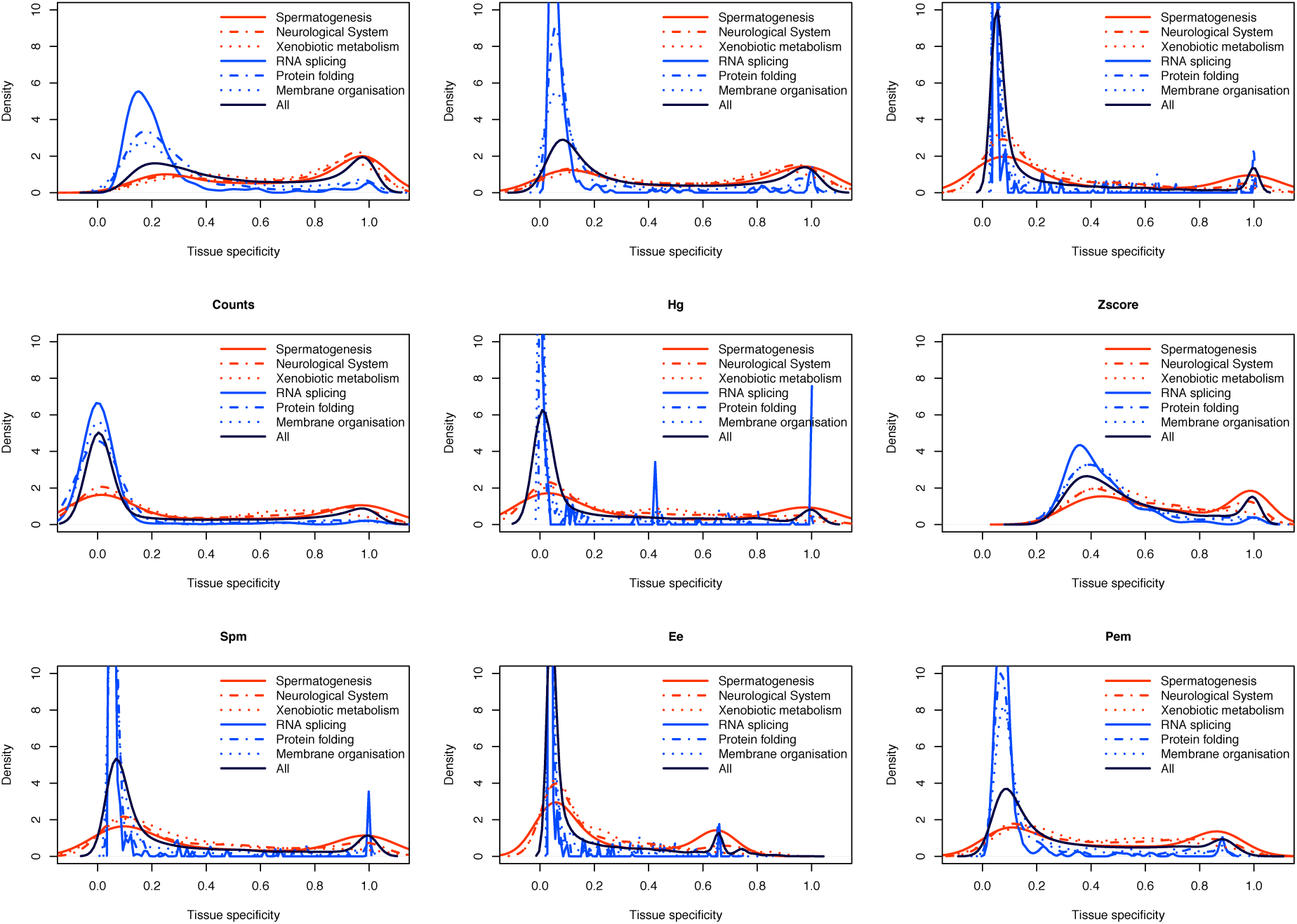
Tissue specificity parameters of subsets of genes which are expected to be tissue-specific (red lines) or broadly expressed (blue lines), based on associated GO terms (described in Material and Methods). The black line represents the distribution for all genes, including those not associated to any of these GO terms.

Most methods seem to have more difficulty in finding tissue-specificity signal than broad expression signal. We checked whether those tissue-specific genes detected by each method are specific to the method, or also detected by others. Strikingly, almost all tissue-specific genes found by any method are also found by Tau. Gini also reports many tissue-specific genes which are reported by Tau but no other method. This is illustrated with the examples of brain and testis specific genes in Figure 5 (for other organs see Figure S17 – S41). To call genes specific a threshold of 0.8 was set, which is after the first peak of the bimodal distribution for most parameters. The same analysis was performed with thresholds of 0.6 and 0.4 (data not shown), and produced similar results: Tau detects all genes that other methods detect plus some that are not detected by any other method. To check whether these additional tissue-specific genes found by Tau are biologically relevant, a GO enrichment test was performed on tissue-specific genes for testis and brain reported by all methods, by Tau alone or only by Tau and Gini (Figure S42 – S47). Each of these genes sets is indeed enriched in brain or testis-specific functions, which shows that these were rather false negatives of the other methods than false positives of Tau and Gini.

**Fig. 5:**
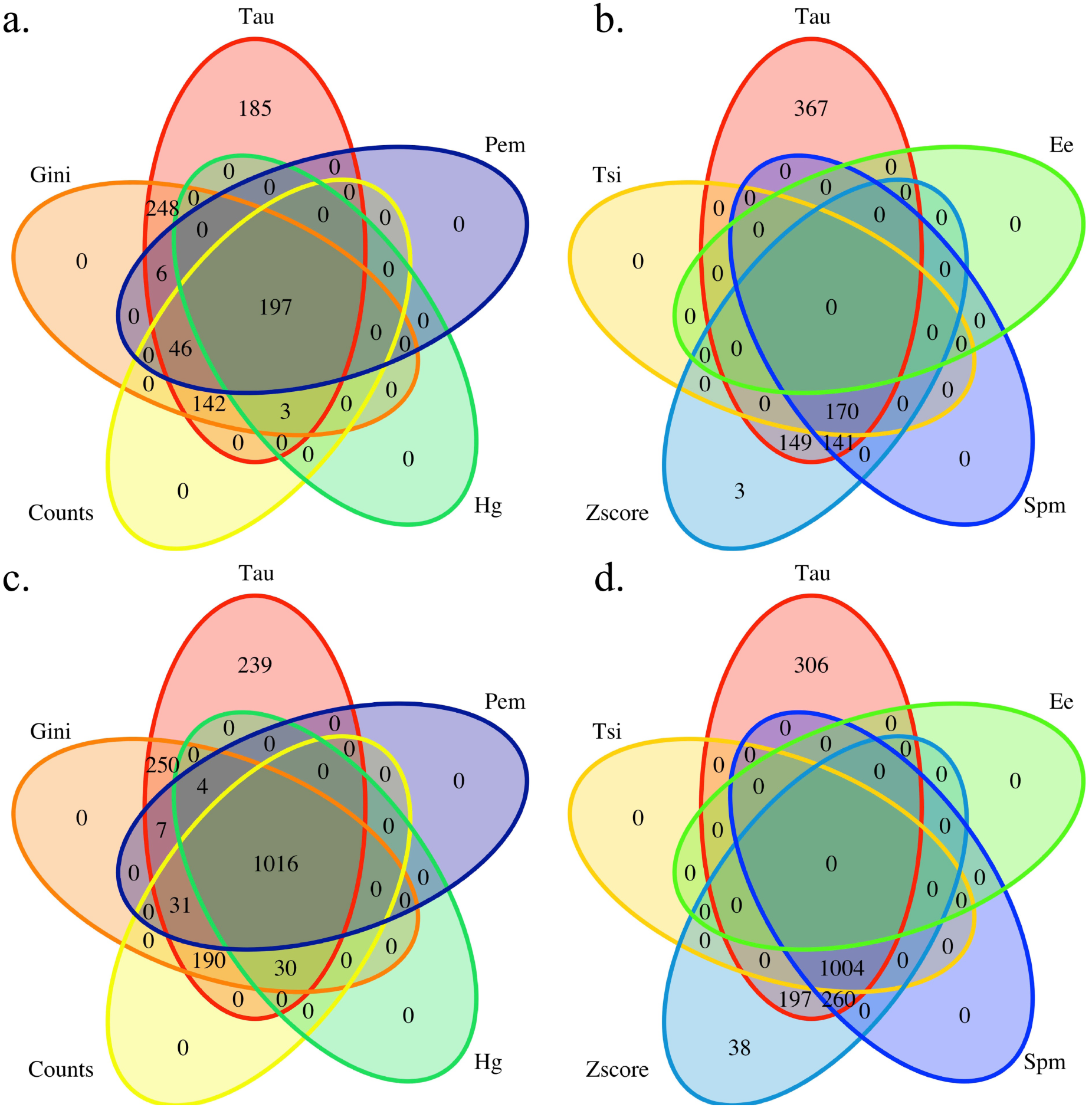
Venn diagram of genes called specific with different parameters, with a cut-off of 0.8 for each parameter; a) and b) genes with their highest expression in the brain; c) and d) genes with their highest expression in the testis; parameters are shown in a/c or b/d for readability, with Tau in common because it calls the most genes tissue-specific.

The same analysis was also performed on the microarray data sets for mouse and human. We compared each metric on a full microarray human data set of 32 tissues and on the subset of 5 tissues (Figure S48). For human, the correlations are weaker than with RNA-seq, even for the best performing metrics: Counts, Gini and Tau (mean 0.2 < r < 0.4). For mouse the correlations on microarray data are better: Counts, Gini and Tau (mean 0.4 < r < 0.6) (Figure S49). Results for 32 and 14 human tissues, and for 19 and 14 mouse tissues, are shown in Figure S50 and S51. The distribution of correlations of Tau calculated on different subsets of tissues is shown in Figure S52 and S53. Similarly, in the comparison between human and mouse orthologs, the correlations are much weaker for microarrays then for RNA-seq (Figure S54). Specificity values are better correlated between RNA-seq and microarray for the mouse then for the human data sets (Figure 6 and S55). This correlation is on the same scale as that between two different RNA-seq datasets, although the correlation is a bit stronger for the RNA-seq datasets (Figure S56 and S57). It should be noted that microarray and RNA-seq can only be compared on the subset of genes for which microarray data is usable, which excludes very tissue-specific genes detected only by RNA-seq (Figure S58).

**Fig. 6:**
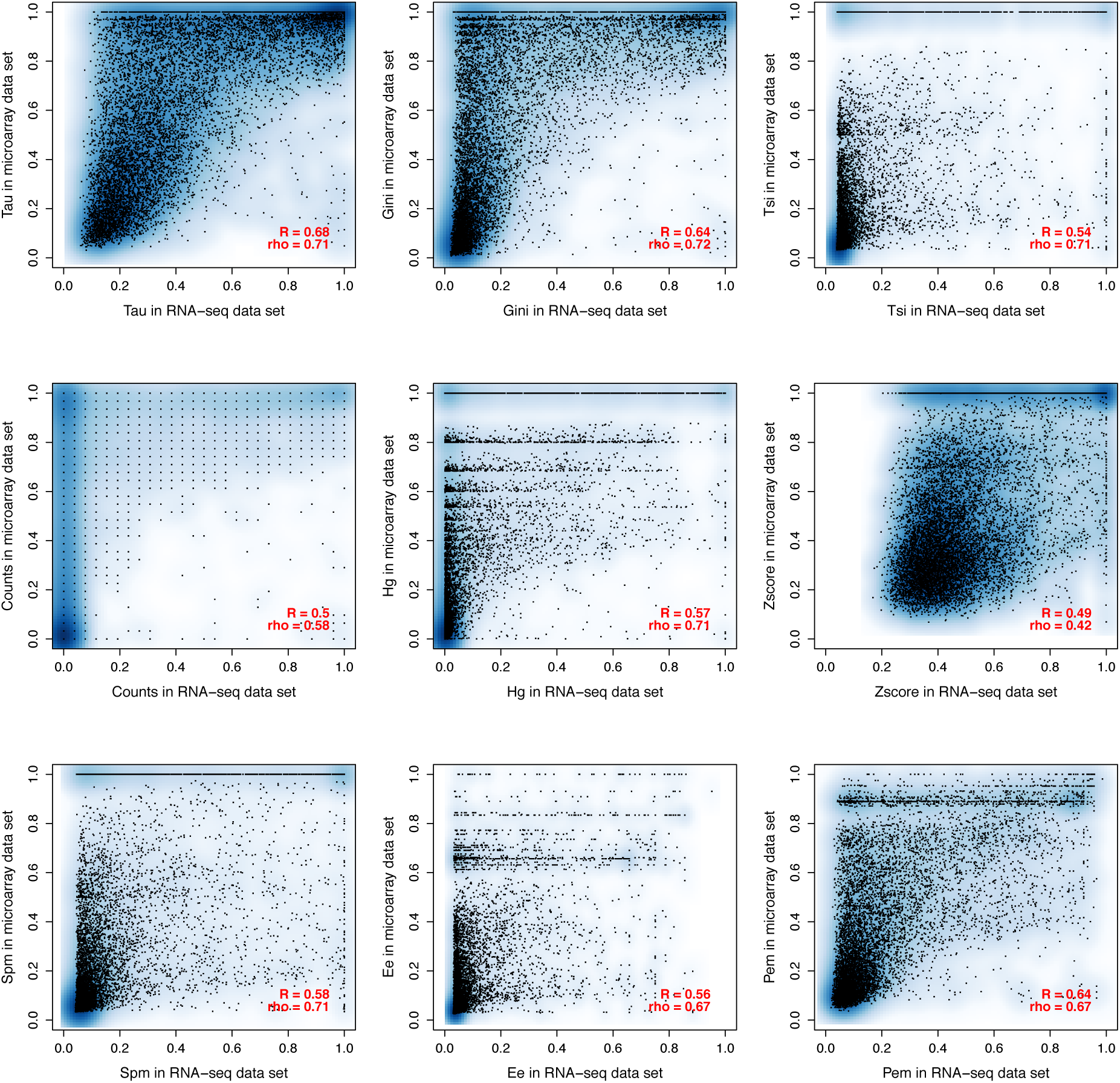
Comparison between tissue specificity parameters calculated on RNA-seq 27 tissues vs. microarray 32 tissues human data sets. All correlations have p-value < 2.2*10^−16^.

Tissue-specificity has been reported to be negatively correlated to mean or maximum gene expression level across tissues, i.e. ubiquitous genes have higher expression and specific genes have lower expression (discussed in (Kryuchkova-Mostacci & Robinson-Rechavi 2015; Subramanian & Kumar 2004; Lercher et al. 2002)). Indeed, we find a negative correlation of all metrics with mean expression; this correlation is similar for RNA-seq (r from −0.69 to − 0.93) and for microarray (r from −0.70 to −0.95) (Figure S59 − S62). Z-score has the weakest correlation with mean expression on RNA-seq data and on microarray data. The correlation of tissue specificity parameters and maximal expression is also similar with RNA-seq and microarray (Figure S63 − S66): all the parameters are negatively correlated with maximal expression.

In all the analyses described above, RPKM values were log-transformed, as described in Material and Methods. In the following we investigated how stable the results of tissue specificity are if data are not log-normalised or if they are additionally quantile normalised. We compared tissue specificity calculated on log-transformed RPKM (as above), raw RPKM, log-transformed and quantile normalised RPKM (Figure S67 − S75). In general quantile normalisation has no influence on the results of calculation of tissue specificity (Figure S74 and S75). Expectedly, removing log-transformation has a greater influence on all parameters, in the direction of detecting more tissue-specificity, sometimes losing completely the signal of broad expression, e.g. Tau (Figure S68). Moreover, in the absence of log-transformation, the correlations between subsets of tissues or between species are in general weaker (Figure S69 − S70). The normalisation has no influence on Counts, as expected, as only yes/no for the expression is taken in the account. Tau, Gini, TSI and Hg show the highest correlations between normalised and non normalised data (Figure S72 and S73), thus appearing more robust.

## Discussion

We analysed nine parameters to calculate tissue specificity. We compared the methods with respect to their stability to number of tissues, their correlation between one-to-one orthologs in human and mouse, their power in detecting tissue-specific genes and their distribution of values. As many experiments do not have many tissues it is important that tissue specificity can be calculated reliably on few tissues.

Different methods of calculating tissue specificity take into account different properties of expression. The Counts method does not take in account the amplitude of differences between tissues. This is the simplest method yet, if the threshold is chosen properly, gives surprisingly good results. Distribution of Counts tissue specificity depending on the chosen threshold is presented in Figure S76: with too high or too low threshold, most genes are reported as not specific, but it is quite robust to a change of one order of magnitude (1 to 10 RPKM). Tau and TSI both use the information of expression of a gene in each tissue and its maximal expression over all tissues. The difference between Tau and TSI is that Tau also takes into account the number of tissues. The Hg coefficient is also similar, but differs in that instead of the maximal expression (necessarily in a specific tissue) the sum of expression over tissues is used; and each normalised value is multiplied by log of the value. And for the SPM score each value (squared) is corrected by the sum of squared gene expression across all tissues. The EE score also corrects each expression value by the sum of gene expression across tissues as well as by the sum of expression in the target tissue. The PEM score is simply the logarithm (base 10) of the EE score. As these coefficients are normalised by either maximal expression of the gene or by the sum of expression of the gene, they are not sensitive to its absolute expression level. Z-score is the only method that takes the standard deviation of expression into account. An overview of the methods with their shared components (e.g., max expression appears in Tau and in TSI) is presented in Supplementary Materials.

Tau appears consistently to be the most robust method in our analyses. Comparing coefficients calculated on different sized data sets, Tau showed one of the highest correlations (Figure 2 and S7-S9). And while it may be debated what is it the “best” distribution between ubiquitous and specific genes, we note that Tau provides well separated groups with lower skew towards calling most genes ubiquitous or tissue-specific than other methods (Figure 1 and S2); and it found more tissue-specific genes (Figure 5, S10, S11 and S16 – S41). Tau also showed a robust behaviour according to normalisation of data (Figure S72 and S73). With the GO analysis performed Tau the best in recognising tissue specific genes (Figure 4 and S15), and conversely tissue-specific genes found only with Tau have functional annotations which are consistent with their tissue of highest expression (Figure S41 – 44).

When a score per tissue is needed, the PEM score showed acceptable results, except for non log-transformed mouse RNA-seq (Figure S73), and it is most similar to Tau. An association between scores and tissues can be also obtained by simply using Tau and choosing the tissue with the highest expression.

Z-score and PEM score are the only methods to detect under-expression. But z-score is very sensitive to the number of tissues used for analysis, and generally performs poorly on most tests. The PEM score performs relatively well, though it is skewed to 0, i.e. to calling genes as ubiquitous (Figure 1 and S2).

In general almost twice as many genes can be called expressed in at least one tissue with RNA-seq than with microarray (see Material and Methods and Figure S58). It has been reported that the detection of lowly expressed genes is better with RNA-seq then with microarrays (Zhao et al. 2014; Wang et al. 2014; Emig et al. 2011). Since the most tissue-specific genes are often lowly expressed (Kryuchkova-Mostacci & Robinson-Rechavi 2015; Subramanian & Kumar 2004; Lercher et al. 2002), RNA-seq can detect specific genes which were not detected using microarrays (Figure S58). We observe that the correlation between RNA-seq and microarray data set is of the same scale as the correlation between two RNA-seq data sets (Figure 6, S55 – S57). It should be noted that the correlation between microarray and RNA-seq is calculated only on half of the genes, mostly excluding specific ones; and that the second RNA-seq data set has only 6 tissues, which could make the correlation between RNA-seq data sets weaker.

Generally, the tissue-specificity estimated from different data types appears to be different. This is notable relative to the number of tissues (Figure 2 compared to S48 and S7 compared to S49): tissue specificity calculated on microarray with a small number of tissues is poorly correlated to that with a larger number in human data, but the opposite is seen for mouse data. The correlation between species is higher for RNA-seq than for microarray (Figure 3 and S54). Our observations imply that past results which relied on microarray data for the evolutionary interpretation of tissue-specificity should be treated with great caution.

With any method of calculation tissue specificity, it should be noted that if the proportion of closely related tissues (e.g. different parts of the brain) in the set of tissues is high, the tissue specificity will be biased. Moreover, usually a large proportion of tissue-specific genes are testis specific, so special care should be taken in comparing data sets with and without testis. Thus, in general during the analysis of tissue specificity care should be taken in sampling the tissues used.

For studying the evolution of gene expression, we show here that tissue-specificity is a biologically relevant parameter which has strong conservation between relatively closely related species such as human and mouse. Our results show that using a robust method such as Tau allows evolutionary comparisons even when tissue sampling somewhat differs (e.g., correlation with 27 vs. 16 tissues). In light of the difficulties of comparing expression levels between species (Piasecka et al. 2012; Pereira et al. 2009; Gilad & Mizrahi-Man 2015), tissue-specificity holds promise not only as a confounding factor to take into account in molecular evolution (Kryuchkova-Mostacci & Robinson-Rechavi 2015), but also as a measure of biological function which can be compared between genes and between species.

## Conclusion

The best overall method to measure expression specificity appears to be Tau, which is reassuring considering the number of studies in which it has been used. Counts is the simplest method, and if the threshold is chosen properly, shows good results, although with a tendency to under-call tissue-specific genes. Gini is very similar to Tau in their performance. These methods allow to capture a signal which has both functional and evolutionary significance to the genes which are studied.

## Acknowledgements

This work was supported by the Swiss National Science Foundation (grants number 31003A 133011/1 and 31003A_153341/1) and Etat de Vaud. We thank Marta Rosikiewicz, Iakov Davydov and Andrea Komljenovic for helpful comments and suggestions.

## References

Assis R, Bachtrog D. 2013. Neofunctionalization of young duplicate genes in Drosophila. PNAS. 110: 17409–14.

Assis R, Bachtrog D. 2015. Rapid divergence and diversification of mammalian duplicate gene functions. BMC Evol. Biol. 15: 138.

Assis R, Kondrashov AS. 2014. Conserved proteins are fragile. Mol. Biol. Evol. 31: 419–24.

Bastian F, Parmentier G, Roux J, Moretti S, Lauder V, et al. 2008. Bgee: integrating and comparing heterogeneous transcriptome data among species. In: Data Integration in the Life Sciences. Springer Berlin Heidelberg pp. 124–131.

Bolstad BM. preprocessCore: A collection of pre-processing functions.

Brawand D, Soumillon M, Necsulea A, Julien P, Csárdi G, et al. 2011. The evolution of gene expression levels in mammalian organs. Nature. 478: 343–8.

Bush SJ, Kover PX, Urrutia AO. 2015. Lineage-specific sequence evolution and exon edge conservation partially explain the relationship of evolutionary rate and expression level in A. thaliana. Mol. Ecol. 24: 3093–3106.

Ceriani L, Verme P. 2012. The origins of the Gini index: Extracts from Variabilità e Mutabilità (1912) by Corrado Gini. J. Econ. Inequal. 10: 421–443.

Chen H. 2015. VennDiagram: Generate High-Resolution Venn and Euler Plots.

Cortez D, Marin R, Toledo-Flores D, Froidevaux L, Liechti A, et al. 2014. Origins and functional evolution of Y chromosomes across mammals. Nature. 508: 488–93.

Cui P, Lin Q, Ding F, Hu S, Yu J. 2012. The transcript-centric mutations in human genomes. Genomics, Proteomics Bioinforma. 10: 11–22.

Dezso Z, Nikolsky Y, Sviridov E, Shi W, Serebriyskaya T, et al. 2008. A comprehensive functional analysis of tissue specificity of human gene expression. BMC Biol. 6: 49.

Divina P, Vlcek C, Strnad P, Paces V, Forejt J. 2005. Global transcriptome analysis of the C57BL/6J mouse testis by SAGE: evidence for nonrandom gene order. BMC Genomics. 6: 29.

Duret L, Mouchiroud D. 2000. Determinants of substitution rates in mammalian genes: expression pattern affects selection intensity but not mutation rate. Mol. Biol. Evol. 17: 68–74.

Eden E, Navon R, Steinfeld I, Lipson D, Yakhini Z. 2009. GOrilla: a tool for discovery and visualization of enriched GO terms in ranked gene lists. BMC Bioinformatics. 10: 48.

Emig D, Kacprowski T, Albrecht M. 2011. Measuring and analyzing tissue specificity of human genes and protein complexes. EURASIP J. Bioinforma. Syst. Biol. 2011: 5.

Fagerberg L, Hallstrom BM, Oksvold P, Kampf C, Djureinovic D, et al. 2013. Analysis of the human tissue-specific expression by genome-wide integration of transcriptomics and antibody-based proteomics. Mol. Cell. Proteomics. 13: 397–406.

Ge X, Yamamoto S, Tsutsumi S, Midorikawa Y, Ihara S, et al. 2005. Interpreting expression profiles of cancers by genome-wide survey of breadth of expression in normal tissues. Genomics. 86: 127–141.

Gilad Y, Mizrahi-Man O. 2015. A reanalysis of mouse ENCODE comparative gene expression data. F1000Research. 121: 1–32.

Handcock MS. 2015. Relative Distribution Methods.

Handcock MS, Morris M. 1999. Relative Distribution Methods in the Social Sciences. Springer: New York.

Hebenstreit D, Fang M, Gu M, Charoensawan V, van Oudenaarden A, et al. 2011. RNA sequencing reveals two major classes of gene expression levels in metazoan cells. Mol. Syst. Biol. 7: 497.

Huminiecki L, Lloyd AT, Wolfe KH. 2003. Congruence of tissue expression profiles from Gene Expression Atlas, SAGEmap and TissueInfo databases. BMC Genomics. 4: 31.

Julien P, Brawand D, Soumillon M, Necsulea A, Liechti A, et al. 2012. Mechanisms and evolutionary patterns of mammalian and avian dosage compensation. PLoS Biol. 10:e1001328.

Kryuchkova-Mostacci N, Robinson-Rechavi M. 2015. Tissue-specific evolution of protein coding genes in human and mouse. PLoS One. 10:e0131673.

Lercher MJ, Urrutia AO, Hurst LD. 2002. Clustering of housekeeping genes provides a unified model of gene order in the human genome. Nat. Genet. 31: 180–3.

Liao B-Y, Scott NM, Zhang J. 2006. Impacts of gene essentiality, expression pattern, and gene compactness on the evolutionary rate of mammalian proteins. Mol. Biol. Evol. 23: 2072–80.

Liao B-Y, Zhang J. 2006. Low rates of expression profile divergence in highly expressed genes and tissue-specific genes during mammalian evolution. Mol. Biol. Evol. 23: 1119–28.

Lin H, Ouyang S, Egan A, Nobuta K, Haas BJ, et al. 2008. Characterization of paralogous protein families in rice. BMC Plant Biol. 8: 18.

Lin S, Lin Y, Nery JR, Urich M a., Breschi A, et al. 2014. Comparison of the transcriptional landscapes between human and mouse tissues. PNAS. 111: 17224–17229.

Liu W -m., Mei R, Ryder TB, Hubbell E, Dee S, et al. 2002. Analysis of high density expressino microarrays with signed-rank call algorithms. Bioinformatics. 18: 1593–1599.

Liu X, Yu X, Zack DJ, Zhu H, Qian J. 2008. TiGER: a database for tissue-specific gene expression and regulation. BMC Bioinformatics. 9: 271.

Ma L, Cui P, Zhu J, Zhang Z, Zhang Z. 2014. Translational selection in human: more pronounced in housekeeping genes. Biol. Direct. 9: 17.

Milnthorpe AT, Soloviev M. 2012. The use of EST expression matrixes for the quality control of gene expression data. PLoS One. 7:e32966.

Park SG, Choi SS. 2010. Expression breadth and expression abundance behave differently in correlations with evolutionary rates. BMC Evol. Biol. 10: 241.

Pereira V, Waxman D, Eyre-Walker A. 2009. A Problem With the Correlation Coefficient as a Measure of Gene Expression Divergence. Genetics. 183: 1597–1600.

Piasecka B, Robinson-Rechavi M, Bergmann S. 2012. Correcting for the bias due to expression specificity improves the estimation of constrained evolution of expression between mouse and human. Bioinformatics. 28: 1865–72.

Ponger L, Duret L, Mouchiroud D. 2001. Determinants of CpG islands: expression in early embryo and isochore structure. Genome Res. 11: 1854–1860.

R Core Team. 2015. R: A language and environment for statistical computing.

Ramsköld D, Wang ET, Burge CB, Sandberg R. 2009. An abundance of ubiquitously expressed genes revealed by tissue transcriptome sequence data. PLoS Comput. Biol. 5:e1000598.

Rosikiewicz M, Robinson-Rechavi M. 2014. IQRray, a new method for Affymetrix microarray quality control, and the homologous organ conservation score, a new benchmark method for quality control metrics. Bioinformatics. 30: 1392–1399.

Russ J, Futschik ME. 2010. Comparison and consolidation of microarray data sets of human tissue expression. BMC Genomics. 11: 305.

Schug J, Schuller W-P, Kappen C, Salbaum JM, Bucan M, et al. 2005. Promoter features related to tissue specificity as measured by Shannon entropy. Genome Biol. 6:R33.

Schuster EF, Blanc E, Partridge L, Thornton JM. 2007. Correcting for sequence biases in present/absent calls. Genome Biol. 8:R125.

Smeds L, Warmuth V, Bolivar P, Uebbing S, Burri R, et al. 2015. Evolutionary analysis of the female-specific avian W chromosome. Nat. Commun. 6: 7330.

Subramanian S, Kumar S. 2004. Gene expression intensity shapes evolutionary rates of the proteins encoded by the vertebrate genome. Genetics. 168: 373–81.

Supek F, Bošnjak M, Škunca N, Šmuc T. 2011. REVIGO Summarizes and Visualizes Long Lists of Gene Ontology Terms. PLoS One. 6:e21800.

The ENCODE Project Consortium. 2011. A user’s guide to the encyclopedia of DNA elements (ENCODE). PLoS Biol. 9:e1001046.

Thorrez L, Van Deun K, Tranchevent LC, Van Lommel L, Engelen K, et al. 2008. Using ribosomal protein genes as reference: A tale of caution. PLoS One. 3:e1854.

Vandenbon A, Nakai K. 2010. Modeling tissue-specific structural patterns in human and mouse promoters. Nucleic Acids Res. 38: 17–25.

Vinogradov AE. 2003. Isochores and tissue-specificity. Nucleic Acids Res. 31: 5212–5220.

Wagner GP, Kin K, Lynch VJ. 2013. A model based criterion for gene expression calls using RNA-seq data. Theory Biosci. 132: 159–164.

Wang C, Gong B, Bushel PR, Thierry-Mieg J, Thierry-Mieg D, et al. 2014. The concordance between RNA-seq and microarray data depends on chemical treatment and transcript abundance. Nat. Biotechnol. 32: 926–32.

Warnes G, Bolker B, Bonebakker L, Gentleman R, Liaw WHA, et al. 2015. gplots: Various R programming tools for plotting data.

Weber CC, Hurst LD. 2011. Support for multiple classes of local expression cluster in Drosophila melanogaster, but no evidence for gene order conservation. Genome Biol. 12:R23.

Winter EE, Goodstadt L, Ponting CP. 2004. Elevated rates of protein secretion , evolution , and disease among tissue-specific genes. Genome Res. 14: 54–61.

Xiao S-J, Zhang C, Zou Q, Ji Z-L. 2010. TiSGeD: a database for tissue-specific genes. Bioinformatics. 26: 1273–5.

Yanai I, Benjamin H, Shmoish M, Chalifa-Caspi V, Shklar M, et al. 2005. Genome-wide midrange transcription profiles reveal expression level relationships in human tissue specification. Bioinformatics. 21: 650–9.

Yu X, Lin J, Zack DJ, Qian J. 2006. Computational analysis of tissue-specific combinatorial gene regulation: predicting interaction between transcription factors in human tissues. Nucleic Acids Res. 34: 4925–36.

Zhao L, Wit J, Svetec N, Begun DJ. 2015. Parallel gene expression differences between low and high latitude populations of Drosophila melanogaster and D. simulans. PLOS Genet. 11:e1005184.

Zhao S, Fung-Leung WP, Bittner A, Ngo K, Liu X. 2014. Comparison of RNA-Seq and microarray in transcriptome profiling of activated T cells. PLoS One. 9:e78644.

